# Prevalence of *Schistosoma mansoni* infection in Ethiopia: A systematic review and meta-analysis

**DOI:** 10.1101/610113

**Authors:** Siraj Hussen, Demissie Assegu, Techalew Shimelis

## Abstract

**Background:** Schistosomiasis is the most predominant helminthic infection in tropics and subtropics mainly in sub-Saharan African countries including Ethiopia. *S. mansoni* infection is still becoming a public health problem since the risk of reinfection and recurrent disease remain, even in areas with high treatment coverage. There is no summarized data regarding prevalence of *S. mansoni* infection in Ethiopia. Therefore, this review was done to determine the pooled prevalence of *S. mansoni* infection in Ethiopia.

**Methods:** The PRISMA guidelines protocol was followed to perform the systematic review and meta-analysis. Published studies from January 1999 to September 1 2018 were searched in Medline, PubMed, Google scholar, HINARI and Cochrane Library. The study search terms were: “prevalence”, “incidence”, “schistosomiasis” “Bilharziasis”, “Ethiopia”. The heterogeneity of studies was assessed using Cochran’s Q test and I^2^ test statistics. Publication bias was assessed by Egger’s test.

**Results:** Eighty four studies were included in this review and meta–analysis. The pooled prevalence of *S. mansoni* among Ethiopian population was 18.7% (95%CI: 14.7-23.5). Southern regions of Ethiopia had a higher *S.mansoni* prevalence of 33.6% 995% CI: 20.2-50.4). *S.mansoni* was higher in rural areas and among males with a pooled prevalence, 20.8% (95% CI: 14.2-29.4) and 29.4% (95%CI: 23.2-36.6), respectively. Similarly, the prevalence of *S.mansoni* have been increased over the past 15 years.

**Conclusion:** The review showed a moderate prevalence of *S.mansoni* infection in Ethiopia and disease is still a major health problem. Therefore, integrated control approach could be implemented to reduce the burden of this parasite in Ethiopia. Interventions leading to reduction of open water sources exposure to reduce schistosomiasis transmission, strengthen of deworming program, giving appropriate health education on the risk of schistosomal infection and transmission should be applied.

**Author Summary:** Understanding summarized data regarding prevalence of *S. mansoni* infection in Ethiopia is essential to inform decisions on appropriate control strategies for schistosomiasis. We searched Published studies from January 1999 to September 1 2018 from Medline, PubMed, Google scholar, HINARI and Cochrane Library. Eighty four studies were included in this review and meta–analysis. The limit of language was English and the limit of study group was human. The pooled prevalence of *S. mansoni* among Ethiopian population was 18.7%. Southern regions of Ethiopia had a higher *S.mansoni* prevalence and the parasite was higher in rural areas and among males. The prevalence of *S.mansoni* have been increased over the past 15 years. Our review showed a moderate prevalence of *S.mansoni* infection in Ethiopia and disease is still a major health problem. Therefore, appropriate controlling approach could be implemented. Interventions leading to reduction of open water sources, strengthen of deworming program, and giving appropriate health education should be applied.

## Background

Schistosomiasis is the most widely distributed chronic but neglected tropical disease (NTD) that affects people living in communities where there is poor environmental sanitation and water supply [1, 2]. Human schistosomiasis is the most deadly NTD and Human schistosomiasis is ranked second to malaria in terms of mortality [1, 2]. An estimated 700 million people in 76 countries are at risk of schistosomiasis, and 240 million people are already infected. About 85% of the infections occur in Africa where a yearly estimated death is 280,000 people and an estimated disability-adjusted life years is 3.3 million people [2–4].

In addition to high morbidity and mortality infection caused by *S. mansoni* among school-age children, adolescents and young adults have the consequences of growth delay and anemia, Vitamin-A deficiency as well as possible cognitive and memory impairment, which limits their potentials in learning [5].

Schistosomiasis is more wide spread in poor rural communities particularly in places where fishing and agricultural activities are dominant. Domestic activities such as washing clothes and fetching water in infected water expose women and children to infection. Poor hygiene and recreational activities like swimming and fishing also increase the risk of infection in children [6, 7].

In Ethiopia, about 5.01 million peoples are infected with schistosomiasis and 37.5 million people are at risk of the parasite [8]. *S. mansoni* is widespread and its presence has been recorded in all administrative regions and is rapidly spreading in connection with water resource development and intensive population movements [9].The optimal altitude category for the transmission of *S. mansoni* is between 1000 and 2000 meters, and most endemic localities in the country are located in this altitudinal range and its prevalence was reported as high as 90% in the country [10, 11]. Two species of fresh water snails (*Biomphalaria pfeifferi* and *Biomphalaria sudanica)* are responsible for the transmission of this parasite in Ethiopia *[12]*.

Reports from the regional mapping survey conducted by the Ethiopian Public Health Institute on schistosomiasis and soil-transmitted helminths across the country showed high distribution of S. mansonia *[13,14]*.The national control programm is designed to achieve elimination for neglected diseases and other poverty related infections, including schistosomiasis as a major public health problem by 2020 and aim to attain transmission break by 2025. Providing a global view of the occurrence of this disease has become a high priority, and rigrous efforts were made to eliminate schistosomiasis through the implementation of sustainable control strategies. However, existing evidence suggests that S. mansoni is still a major public health problem causing significant morbidity and mortality in endemic countries, particularly in Ethiopia. In this study, we used data published from Ethiopia between 1999 and 2018 to perform a systematic review and meta-analysis of the prevalence of S. mansoni to provide information that will help in tackling the disease at the national level..

## Methods

### Search strategy

A comprehensive literature search was conducted from biomedical data bases: Medline, PubMed, Google scholar, HINARI and Cochrane Library using a special index search terms (medical subject headings (MeSH) “prevalence”,“incidence”, “schistosomiasis” “Bilharziasis”, “Ethiopia”, title and abstract. The limit of language was English and the limit of study group was human. Search was carried out for articles published from 1999 to 2018. Age group categorization was done as follows; children were designated as those of 14 years of age and below; adolescent 15– 17 years; and adult 18 years or higher. The Preferred Reporting Items for Systematic Reviews and Meta-analyses (PRISMA) guideline was used to report the result of this systematic review and meta-analyses (Table S1).

#### Selection criteria

Abstracts retrieved from the initial search were screened using defined inclusion and exclusion criteria.

#### Inclusion criteria and Exclusion Criteria

Studies were selected for systematic review and meta-analysis if: 1) they were conducted in Ethiopia, 2) study design was cross-sectional, 3) studies reported the prevalence of *S.mansoni*, 4) studies reported data in humans and were published in the English language.

Studies were examined for eligibility by reading their titles and abstracts. Relevant abstracts were further assessed for inclusion in the list of full text articles. During the article selection process, studies which did not have full texts were excluded since it was not possible to assess the quality of each article in the absence of their full texts.

### Data extraction

The data extraction was done by three researchers (S.H T.S and D.A) using a standardized and pretested format. The data abstraction format included first author, study design, region in Ethiopia, publication year, sample size, study population, number who tested positive and prevalence of *S.mansoni*. Disagreement on data extractions between researchers was resolved through discussion and consensus.

### Quality assessment

The quality of each article was assessed using 9 point Joanna Briggs Institute (JBI) critical appraisal tools. The tool uses the following criteria: 1) sample frame appropriate to address the target population, 2) study participants sampled in an appropriate way, 3) adequate sample size, 4) study participants sampled in an appropriate way, 5) study subjects and the setting described in detail, 6) data analysis conducted with sufficient coverage of the identified sample, 7) valid methods used for the identification of the condition and the condition was measured in a standard and reliable way for all participants, 8) appropriate statistical analysis; and, 9) adequate response rate. Individual studies were assigned a score that was computed using different parameters in line with the review objectives. The responses were scored 0 for “Not reported” and 1 for “Yes”. Total scores ranged between 0 and 9. Studies with medium (fulfilling 50% of quality assessment parameter) and high quality were included for analysis [15]. None of the studies were excluded based on the quality assessment criteria (Additional file S2).

### Statistical analysis

Data entry and analysis were done using Comprehensive Meta-analysis (version 3.1). The summary of pooled prevalence of *S. mansoni* infection with 95% CI was obtained using the random effects model, due to the possibility of heterogeneity among the studies.

### Sub-group analysis

Sub-group analysis was performed based on geographical region; (Amhara, Oromia, Southern Ethiopia, Tigray, Harari and Afar), Year of study; (1999-2003, 2004-2008, 2009-2013, and 2014-2018), laboratory dagnostic test: (Kato-Katz, Kato-Katz & wet mount, wet mount & formol-ethe, Formol-ether, Kato-Katz & formol-ether, Kato-Katz &SAF and wet mount),age groups: (all age groups, children, Children & adolescent, adolescent &adult and Adult), sex (male and female), and study setting (Rural and Urban).

### Heterogeneity and publication bias

Statistical heterogeneity was assessed by Cochran’s Q test, which indicated the amount of heterogeneity between studies and I^2^ statistic. The I^2^ offers an estimate percentage of the variability in effect estimates, that is due to heterogeneity rather than sampling error or chance differences. Therefore, the existence of heterogeneity was confirmed using Cochran’s Q test (P < 0.10 shows statistically significant heterogeneity) [16]. And I^2^ test that measures level of statistical heterogeneity between studies (values of 25 %, 50 % and 75 % are low, medium and high heterogeneity, respectively) [17]. The Egger weighted regression test methods was used to statistically assess publication bias (P<0.05) [18].

## Results

### Identified studies

A total of 140 records were retrieved through electronic database searching. A total of 42 articles were excluded using their title and abstract review. Ninety eight articles were assessed for eligibility and 14 articles were excluded (eight articles are not cross-sectional study and six have no prevalence data). Finally, 84 studies were found to be eligible and were included in the meta– analysis (figure1).

**Figure 1:**
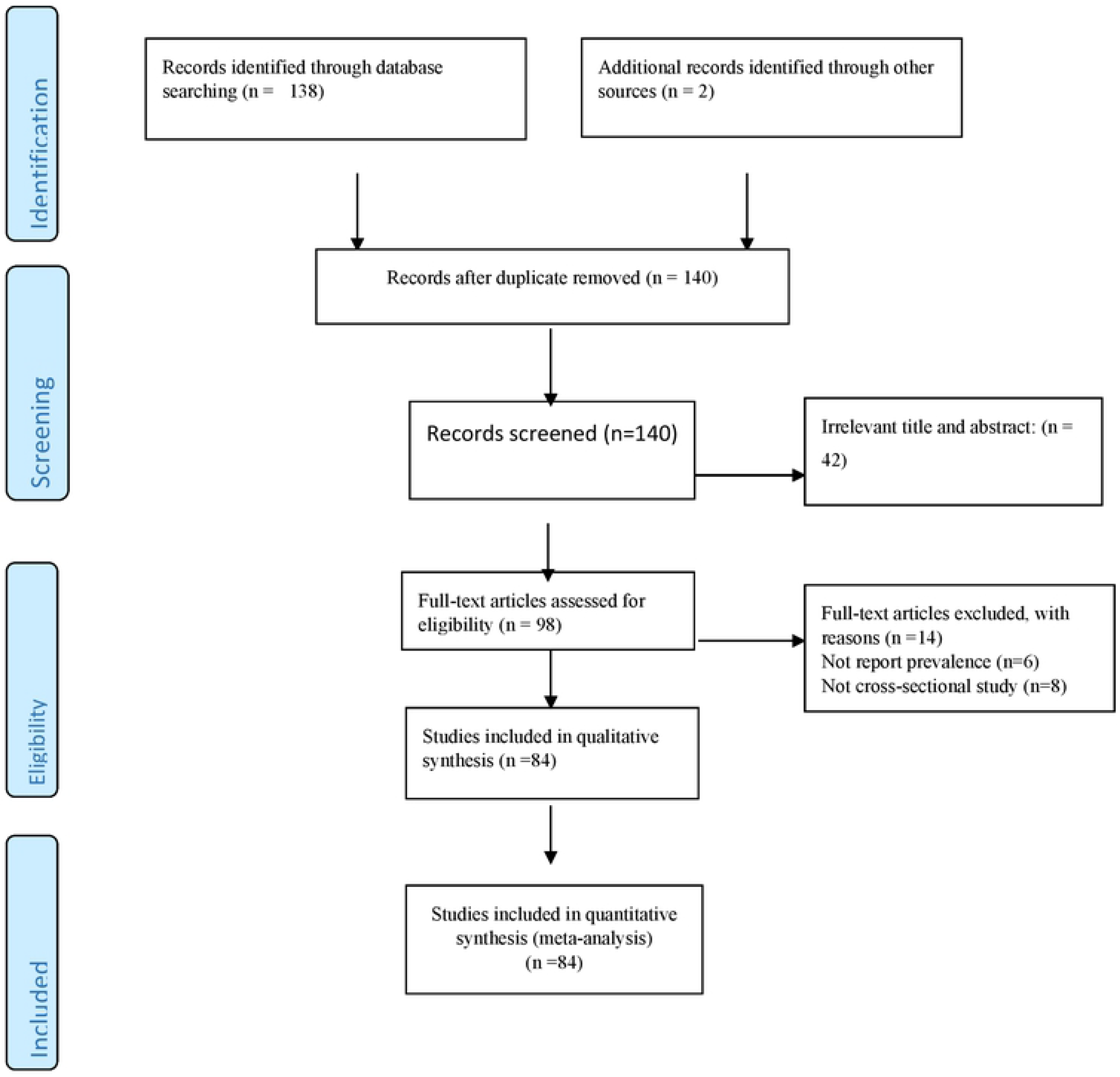
Flow diagram of the studies included in the Meta-analysis

**Figure 2:**
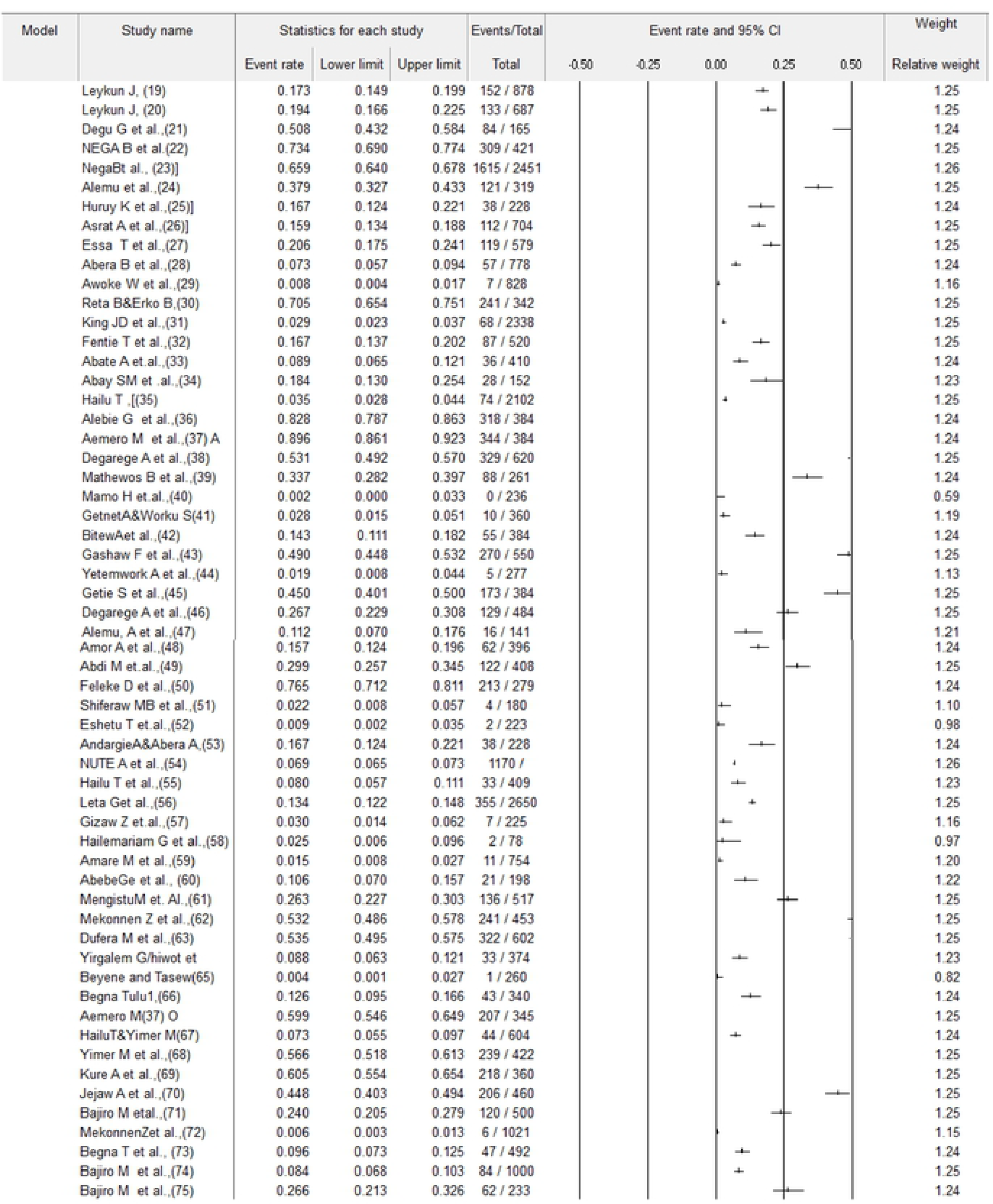
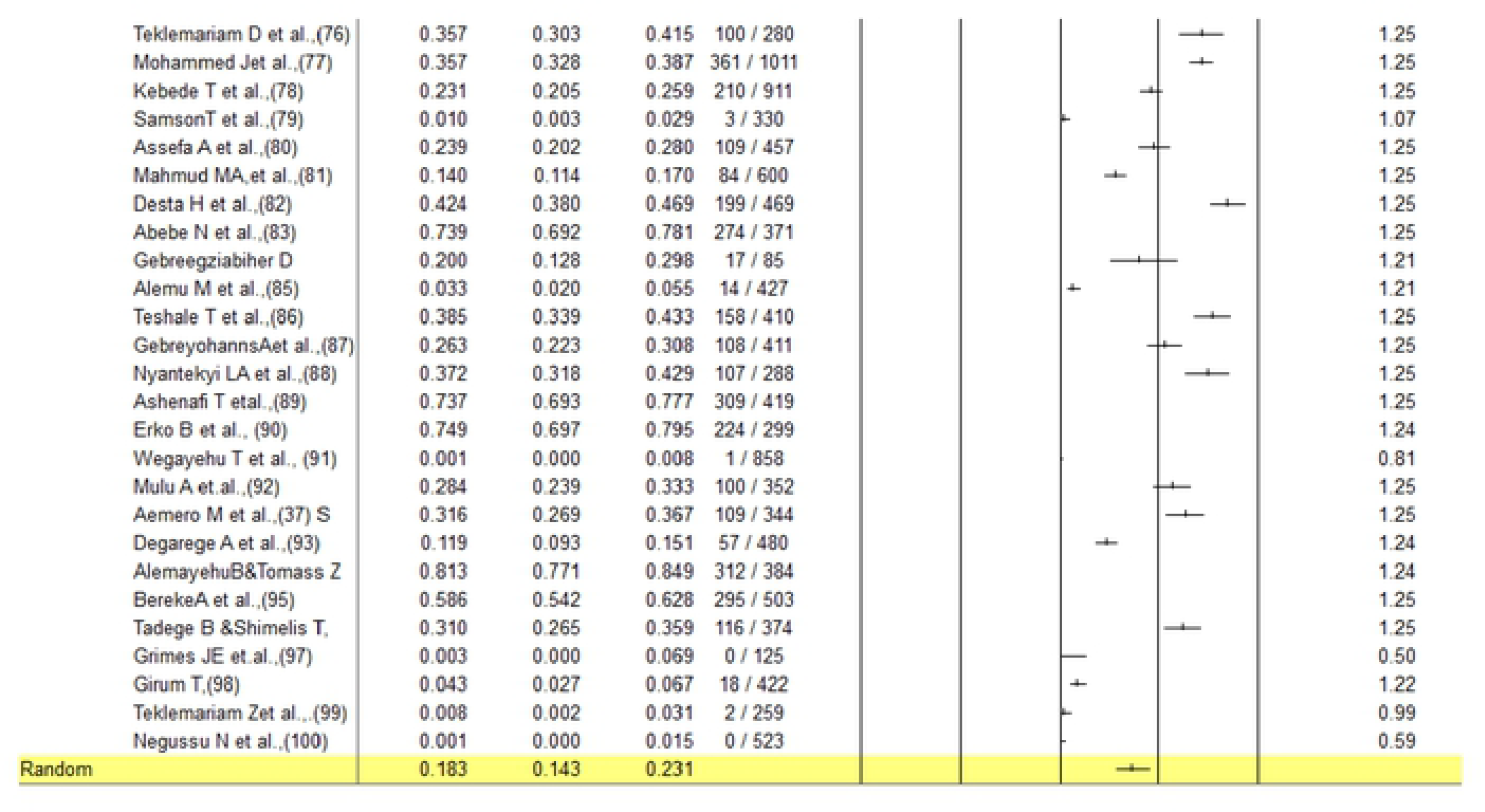
Forest plot for the prevalence of Schistosoma mansoni in Ethiopia

### Study characterstics

In this systematic review and meta-analysis, a total of 60,725 study population was screened for *S. mansoni* infection. Geographically, the population screened for *S. mansoni* infection in six administrative regions of Ethiopia: Amhara, Oromia, Southern Ethiopia, Tigray Harar and Afar (Table 1). The sample size of study population varied from 85 to 16,955 participants (Table 1). The pooled prevalence of *S. mansoni* infection among Ethiopian population was 18.3% (95%CI: 14.3-23.1) (figure 1). There was a high level of heterogeneity, random effect model methods (I^2^ = 99.29, p < 0.001); however, no evidence of publication bias was shown with Egger’s regression intercept (p= 0.639) (s1 figure). The symmetry of funnel plot shows a small publication bias and insignificant effect as portrayed graphically (figure S2). Studies included in this systematic review and meta-analysis were published from 1999 to 2018 and reported from six regions of Ethiopia. The highest and lowest prevalence of *S. mansoni* infection was reported in Amhara (89.6%) and Southern Ethiopia (0.12%), respectively (Table 1).

### Subgroup analysis

Subgroup analysis revealed a broad inconsistency in the prevalence of *S. mansoni* infection among the different parameters used (table 2). By the year of publications, prevalence was highest in 2014 to 2018 publications at 20.9% (95% CI, 15.5-27.5), and the least prevalence was in studies published between 2004-2008 (14.5%, 95% CI, 3.7-42.7). Prevalence was higher in the southern region of Ethiopia (33.6%, 95% CI: 20.2-50.4) (Table 2). Prevalence was also higher in rural areas 20.8% (95% CI: 14.2-29.4) than urban areas 14.9% (95% CI: 9.5-22.8).

The subgroup analysis was done on the prevalence *S.mansoni* by type of diagnostic test. Pooled prevalence of *S.mansoni* was 33.3% (95%CI: 26.5-40.8) using Kato-Katz, 7.4% (95%CI: 2.1-23.0) using Kato-Katz & wet mount, 5.1% (95%CI: 2.5-10.0) using wet mount & formol-ether, 10.7% (95%CI: 6.1-18.2) using Formol-ether, 25.6% (95%CI: 14.3-41.5) using Kato-Katz & formol-ether, 19.4% (95%CI: 0.9-86.4) using Kato-Katz &SAF, and 3.2% (95%CI: 1.7-6.2) using wet mount (Table 2). While grouping the pooled prevalence by sex: 29.4% (95%CI: 23.2-36.6) among males and 22.4% (95%CI: 17.3-28.5) in females.

In this review prevalence of *S. mansoni* was highest in children and adolescents (18.1% (95% CI: 10.3-29.9)), compared to children 10.9% (95% CI: 4.3-25.1) and adolescent & adult 0.5% (95% CI: 1.1-19.8) (Table 2). Meta-regression analysis showed that prevalence of *S. mansoni* had significant association with geographic region (p= 0.0035), study setting (p=0.0242), and with laboratory diagnostic technique (p=0.0001) (figure S3).

## Discussion

Efforts to reduce the epidemiological and clinical consequences of this parasitic infection through the deworming program of the Ethiopian Enhanced Outreach Strategy targeting children below five years of age has been in progress since 2010. However, the burden of S. mansoni infection still remains a public health problem since the risk of reinfection and recurrent disease still exist even in areas with high treatment coverage in the country. We recommend the World Health Organisation’s new focus on transmission control which involves the examination of the efficacy of snail intermediate host control for the prevention of human-snail-human parasite transmission. Different studies have been carried out in different parts of Ethiopia at different times to document the epidemiology of *S. mansoni* infection. However, there is no summarized prevalence data of this parasitic infection at country level to help in the formulation of appropriate intervention methods. Therefore, the present study is the first of its kind and aimed to determine the pooled prevalence of *S. mansoni* in Ethiopia.

In this study, the pooled prevalence of Schistosoma mansoni among Ethiopian population was 18.3% (95%CI: 14.3-23.1). This shows an endemicity and moderate prevalence of *S. mansoni* infection found in Ethiopia [101]. This is comparable with meta-analysis studies conducted in Brazil [102]. However, this finding is higher than the pooled stool S *S.mansoni* estimated from migrants [103]. The difference in prevalence may be due to the different in geographical and ecological variations, periodical cleaning of the irrigation canals, long time endemicity of study area, study design, sampling techniques, sample size, behavior of the study participants, environmental sanitation, and distribution of snails.

Southern Ethiopia had the highest regional prevalence (33.6%), followed by Tgray (20.3%) and then Oromia (18.2%). The variations across regions may be explained by the differences in environmental conditions such as temperature and humidity, rainfall patterns and environmental sanitation which influence parasite transmission. Other factors may include availability and abundance of snail intermediate hosts, socioeconomic conditions, levels of community awareness of the disease, variations in study period, and methods of diagnosis among others. In-spite of the efforts in place towards the control of Schistosomiasis in Ethiopia, our study revealed an increase in the prevalence of the disease during the last 4 years (2014-2018) reviewed probably due to recent water resources development for irrigation and intensive human migration [104]. Furthermore, climate change and global warming which usually result in increased temperature may be additional factors [105]. For instance, a study from Nigeria showed that a rise in ambient temperature from 20-30 OC will lead to an increase in the mean burden of *S. mansoni* [106].

In this study, the pooled prevalence of *S. mansoni* infection in rural settings was higher than that reported from urban settings. This concurs with the report of a systematic review from Kinshasha, Kongo [107]. The higher prevalence from rural settings may be due to increased exposure to water through different activities such as high irrigation practice, swimming and fishing, limited access to health-care services and lack of safe water for the rural population. The limitations of this study were sample size variations, inconsistency of laboratory diagnostic methods used by the individual studies, study periods and regional heterogeneities.

Another important observation was that pooled prevalence of *S. mansoni* is more prevalent in males than females (29.4% versus 22.4%), respectively. This is in agreement with single previous prevalence studies conducted in Ethiopia [(108,109]. The difference in infection rate might be due, males are mostly participated in outdoor activities like irrigation, farming and culturally males exercise swimming and bathing in river water and this may lead to infection by *S.mansoni* cercariae.

Higher pooled prevalence of *S.mansoni* was reported by Kato-Katz tests. Pooled prevalence of *S.mansoni* which used wet mount, wet mount & formol-ether for the diagnosis was low. This could be possibly explained by the high sensitivity of Kato-Katz test for the diagnosis of *S.mansoni* infection. This is in line with WHO 2002 report that have high sensitivity of kato-katz test with a high sensitivity when infection intensity is high in community [110].

The prevalence of *S.mansoni* infection rate was high in children and adolescent than adolescent and adult or adults. This could be associated with children and adolescents are part takers in swimming and recreation. Similar results was reported in in review conducted in Nigeria [11].

### Limitation

The current review and meta-analysis used data which are over-representative of urban populations with greater access to *S.mansoni* prevention and treatment services than rural populations and may underestimate the true burden of this disease in the rural community. Moreover, most of studies which were included in the analysis were clinic/hospital-based studies, and the data might not be representative of the population/community-based prevalence of *S.mansoni* infection. Further, sample size variations, inconsistency of the laboratory diagnostic methods used in the studies as well as study time and regional heterogeneity may affect review of the study.

#### Conclusion and Recommendation

The review showed a moderate prevalence of *S. mansoni* infection in Ethiopians and the diseases is still a major health problem. Therefore, integrated control approach could be implemented to reduce the burden of *S. mansoni* in Ethiopia. Interventions leading to reduction of open water sources exposure to reduce schistosomiasis transmission, strengthening of deworming programs, giving appropriate health education on the risk of schistosomal infection and transmission are suggested.

## Figure legends

Figure S1: Egger regression intercept for the prevalence of Schistosoma mansoni in Ethiopia

Figure S2: Funnel plot for the prevalence of Schistosoma mansoni in Ethiopia

Figure S3: Met regression analysis for the prevalence of Schistosoma mansoni in Ethiopia

## References

1. Organization WHO. Helminth control in school-age children: a guide for managers of control programmes: Geneva: World Health Organization; 2011.

2. Randjelovic A, Frønæs S, Munsami M, Kvalsvig J, Zulu S, Gagai S, et al. A study of hurdles in mass treatment of Schistosomiasis in Kwazulu-Natal, South Africa. South African Family Practice. 2015;57(2):57–61.

3. Adisa J, Egbujo E, Yahaya B, Echejoh G. Primary infertility associated with Schitosoma mansoni: a case report from the Jos plateau, north central Nigeria. African health sciences. 2012;12(4):563–5.

4. Hotez PJ, Alvarado M, Basáñez M-G, Bolliger I, Bourne R, Boussinesq M, et al. The global burden of disease study 2010: interpretation and implications for the Neglected Tropical Diseases. PLoS neglected tropical diseases. 2014;8(7):e2865.

5. Karunamoorthi K, Almalki MJ, Ghailan KY. Schistosomiasis: A neglected tropical disease of poverty: A call for intersectoral mitigation strategies for better health. Journal of Health Research and Reviews. 2018;5(1):1.

6. Organization WH. Schistosomiasis. 2014.

7. Organization WH. Prevention and control of schistosomiasis and soil-transmitted helminthiasis: report of a WHO expert committee. 2002.

8. Steinmann P, Keiser J, Bos R, Tanner M, Utzinger J. Schistosomiasis and water resources development: systematic review, meta-analysis, and estimates of people at risk. The Lancet infectious diseases. 2006;6(7):411–25.

9. Shibru T. Schistosomiasis and its distribution in Ethiopia and Eritrea in: Hailu B, Shibru T, Leykun J, editors. Schistosomiasis in Ethiopia and Eritria 2nd ed Addis Ababa: Institute of Pathobiology Addis Ababa University. 1998:1–18.

10. Zein Ahmed Z, Kloos H. Ecology of health and disease in Ethiopia. 1988.

11. Kabatereine NB, Brooker S, Tukahebwa EM, Kazibwe F, Onapa AW. Epidemiology and geography of Schistosoma mansoni in Uganda: implications for planning control. Tropical Medicine & International Health. 2004;9(3):372–80.

12. Savioli L, Daumerie D. Sustaining the drive to overcome the global impact of neglected tropical diseases: second WHO report on neglected tropical diseases: World Health Organization; 2013.

13. Chitsulo L, Engels D, Montresor A, Savioli L. The global status of schistosomiasis and its control. Acta tropica. 2000;77(1):41–51.

14. Negussu N, Mengistu B, Kebede B, Deribe K, Ejigu E, Tadesse G, et al. Ethiopia schistosomiasis and soil-transmitted helminthes control programme: progress and prospects. Ethiopian medical journal. 2017;55(Suppl 1):75.

15. JBI. critical appraisal checklist for studies reporting prevalence data 2016.

16. DerSimonian R, Laird N. Meta-analysis in clinical trials. Controlled clinical trials. 1986;7(3):177–88.

17. Rücker G, Schwarzer G, Carpenter JR, Schumacher M. Undue reliance on I 2 in assessing heterogeneity may mislead. BMC medical research methodology. 2008;8(1):79.

18. Ioannidis JP. Interpretation of tests of heterogeneity and bias in meta-analysis. Journal of evaluation in clinical practice. 2008;14(5):951–7.

19. Jemaneh L. Intestinal helminth infections in schoolchildren in Gonder town and surrounding areas, Northwest Ethiopia. SINET: Ethiopian Journal of Science. 1999;22(2):209–20.

20. Jemaneh L. Soil-transmitted helminth infections and Schistosomiasis mansoni in school children from Chilga District, Northwest Ethiopia. Ethiopian journal of health sciences. 2001;11(2).

21. Degu G, Mengistu G, Jones J. Praziquantel efficacy against Schistosomia mansoni in schoolchildren in north-west Ethiopia. Transactions of the Royal Society of Tropical Medicine and Hygiene. 2002;96(4):444–5.

22. Berhe N, Halvorsen BL, Gundersen TE, Myrvang B, Gundersen SG, Blomhoff R. Reduced serum concentrations of retinol and α-tocopherol and high concentrations of hydroperoxides are associated with community levels of S. mansoni infection and schistosomal periportal fibrosis in Ethiopian school children. The American journal of tropical medicine and hygiene. 2007;76(5):943–9.

23. Berhe N, Myrvang B, Gundersen SG. Intensity of Schistosoma mansoni, hepatitis B, age, and sex predict levels of hepatic periportal thickening/fibrosis (PPT/F): a large-scale community-based study in Ethiopia. The American journal of tropical medicine and hygiene. 2007;77(6):1079–86.

24. Alemu A, Atnafu A, Addis Z, Shiferaw Y, Teklu T, Mathewos B, et al. Soil transmitted helminths and Schistosoma mansoni infections among school children in Zarima town, northwest Ethiopia. BMC infectious diseases. 2011;11(1):189.

25. Huruy K, Kassu A, Mulu A, Worku N, Fetene T, Gebretsadik S, et al. Intestinal parasitosis and shigellosis among diarrheal patients in Gondar teaching hospital, northwest Ethiopia. BMC research notes. 2011;4(1):472.

26. Ayalew A, Debebe T, Worku A. Prevalence and risk factors of intestinal parasites among Delgi school children, North Gondar, Ethiopia. Journal of Parasitology and Vector Biology. 2011;3(5):75–81.

27. Essa T, Birhane Y, Endris M, Moges A, Moges F. Current status of Schistosoma mansoni infections and associated risk factors among students in Gorgora town, Northwest Ethiopia. ISRN Infectious Diseases. 2012;2013.

28. Abera B, Alem G, Yimer M, Herrador Z. Epidemiology of soil-transmitted helminths, Schistosoma mansoni, and haematocrit values among schoolchildren in Ethiopia. The Journal of Infection in Developing Countries. 2013;7(03):253–60.

29. Awoke W, Bedimo M, Tarekegn M. Prevalence of schistosomiasis and associated factors among students attending at elementary schools in Amibera District, Ethiopia. Open Journal of Preventive Medicine. 2013;3(02):199.

30. Reta B, Erko B. Efficacy and side effects of praziquantel in the treatment for Schistosoma mansoni infection in school children in Senbete Town, northeastern E thiopia. Tropical Medicine & International Health. 2013;18(11):1338–43.

31. King JD, Endeshaw T, Escher E, Alemtaye G, Melaku S, Gelaye W, et al. Intestinal parasite prevalence in an area of Ethiopia after implementing the SAFE strategy, enhanced outreach services, and health extension program. PLoS neglected tropical diseases. 2013;7(6):e2223.

32. Fentie T, Erqou S, Gedefaw M, Desta A. Epidemiology of human fascioliasis and intestinal parasitosis among schoolchildren in Lake Tana Basin, northwest Ethiopia. Transactions of the Royal Society of Tropical Medicine and Hygiene. 2013;107(8):480–6.

33. Abate A, Kibret B, Bekalu E, Abera S, Teklu T, Yalew A, et al. Cross-sectional study on the prevalence of intestinal parasites and associated risk factors in Teda Health Centre, Northwest Ethiopia. ISRN parasitology. 2013;2013.

34. Abay SM, Tilahun M, Fikrie N, Habtewold A. Plasmodium falciparum and Schistosoma mansoni coinfection and the side benefit of artemether-lumefantrine in malaria patients. The Journal of Infection in Developing Countries. 2013;7(06):468–74.

35. Hailu T. Current prevalence of intestinal parasites emphasis on hookworm and schistosoma mansoni infections among patients at Workemeda Health Center, Northwest Ethiopia. Clinical Microbiology: Open Access. 2014.

36. Alebie G, Erko B, Aemero M, Petros B. Epidemiological study on Schistosoma mansoni infection in Sanja area, Amhara region, Ethiopia. Parasites & vectors. 2014;7(1):15.

37. Aemero M, Berhe N, Erko B. Status of Schistosoma mansoni prevalence and intensity of infection in geographically apart endemic localities of Ethiopia: A comparison. Ethiopian journal of health sciences. 2014;24(3):189–94.

38. Degarege A, Legesse M, Medhin G, Teklehaymanot T, Erko B. Day-to-day fluctuation of point-of-care circulating cathodic antigen test scores and faecal egg counts in children infected with Schistosoma mansoni in Ethiopia. BMC infectious diseases. 2014;14(1):210.

39. Mathewos B, Alemu A, Woldeyohannes D, Alemu A, Addis Z, Tiruneh M, et al. Current status of soil transmitted helminths and Schistosoma mansoni infection among children in two primary schools in North Gondar, Northwest Ethiopia: a cross sectional study. BMC Research Notes. 2014;7(1):88.

40. Mamo H. Intestinal parasitic infections among prison inmates and tobacco farm workers in Shewa Robit, north-central Ethiopia. PLoS One. 2014;9(6):e99559.

41. Getnet A, Worku S. The Association Between Major Helminth Infections (Soil-Transmitted Helminthes and Schistosomiasis) and Anemia Among School Children in Shimbit Elementary School, Bahir Dar, Northwest Ethiopia. American Journal of Health Research. 2015;3(2):97–104.

42. Bitew AA, Abera B, Seyoum W, Endale B, Kiber T, Goshu G, et al. Soil-transmitted helminths and Schistosoma mansoni infections in Ethiopian Orthodox Church students around lake tana, northwest Ethiopia. PloS one. 2016;11(5):e0155915.

43. Gashaw F, Aemero M, Legesse M, Petros B, Teklehaimanot T, Medhin G, et al. Prevalence of intestinal helminth infection among school children in Maksegnit and Enfranz Towns, northwestern Ethiopia, with emphasis on Schistosoma mansoni infection. Parasites & vectors. 2015;8(1):567.

44. Aleka Y, Tamir W, Birhane M, Alemu A. Prevalence and Associated Risk Factors of Intestinal Parasitic Infection among Underfive Children in University of Gondar Hospital, Gondar, Northwest Ethiopia. Biomedical Research and Therapy. 2015;2(08):347–53.

45. Getie S, Wondimeneh Y, Getnet G, Workineh M, Worku L, Kassu A, et al. Prevalence and clinical correlates of Schistosoma mansoni co-infection among malaria infected patients, Northwest Ethiopia. BMC research notes. 2015;8(1):480.

46. Degarege A, Hailemeskel E, Erko B. Age-related factors influencing the occurrence of undernutrition in northeastern Ethiopia. BMC public health. 2015;15(1):108.

47. Alemu A, Tegegne Y, Damte D, Melku M. Schistosoma mansoni and soil-transmitted helminths among preschool-aged children in Chuahit, Dembia district, Northwest Ethiopia: prevalence, intensity of infection and associated risk factors. BMC public health. 2016;16(1):422.

48. Amor A, Rodriguez E, Saugar JM, Arroyo A, López-Quintana B, Abera B, et al. High prevalence of Strongyloides stercoralis in school-aged children in a rural highland of north-western Ethiopia: the role of intensive diagnostic work-up. Parasites & vectors. 2016;9(1):617.

49. Abdi M, Nibret E, Munshea A. Prevalence of intestinal helminthic infections and malnutrition among schoolchildren of the Zegie Peninsula, northwestern Ethiopia. Journal of infection and public health. 2017;10(1):84–92.

50. Feleke DG, Arega S, Tekleweini M, Kindie K, Gedefie A. Schistosoma mansoni and other helminthes infections at Haike primary school children, North-East, Ethiopia: a cross-sectional study. BMC research notes. 2017;10(1):609.

51. Shiferaw MB, Zegeye AM, Mengistu AD. Helminth infections and practice of prevention and control measures among pregnant women attending antenatal care at Anbesame health center, Northwest Ethiopia. BMC research notes. 2017;10(1):274.

52. Eshetu T, Sibhatu G, Megiso M, Abere A, Baynes HW, Biadgo B, et al. Intestinal Parasitosis and Their Associated Factors among People Living with HIV at University of Gondar Hospital, Northwest-Ethiopia. Ethiopian journal of health sciences. 2017;27(4):411–20.

53. Andargie AA, Abera AS. Determinants of Schistosoma mansoni in Sanja health center, north West Ethiopia. BMC public health. 2018;18(1):620.

54. Nute AW, Endeshaw T, Stewart AE, Sata E, Bayissasse B, Zerihun M, et al. Prevalence of soil-transmitted helminths and Schistosoma mansoni among a population-based sample of school-age children in Amhara region, Ethiopia. Parasites & vectors. 2018;11(1):431.

55. Hailu T, Alemu M, Abera B, Mulu W, Yizengaw E, Genanew A, et al. Multivariate analysis of factors associated with Schistosoma mansoni and hookworm infection among primary school children in rural Bahir Dar, Northwest Ethiopia. Tropical diseases, travel medicine and vaccines. 2018;4(1):4.

56. Leta GT, French M, Dorny P, Vercruysse J, Levecke B. Comparison of individual and pooled diagnostic examination strategies during the national mapping of soil-transmitted helminths and Schistosoma mansoni in Ethiopia. PLoS neglected tropical diseases. 2018;12(9):e0006723.

57. Gizaw Z, Adane T, Azanaw J, Addisu A, Haile D. Childhood intestinal parasitic infection and sanitation predictors in rural Dembiya, northwest Ethiopia. Environmental Health and Preventive Medicine. 2018;23(1):26.

58. Hailemariam G, Kassu A, Abebe G, Abate E, Damte D, Mekonnen E, et al. Intestinal parasitic infections in HIV/AIDS and HIV seronegative individuals in a teaching hospital, Ethiopia. Japanese journal of infectious diseases. 2004;57(2):41–3.

59. Mengistu A, Gebre-Selassie S, Kassa T. Prevalence of intestinal parasitic infections among urban dwellers in southwest Ethiopia. Ethiopian Journal of Health Development. 2007;21(1):12–7.

60. Abebe G, Kiros M, Golasa L, Zeynudin A. Schistosoma mansoni infection among patients visiting a health centre near Gilgel Gibe Dam, Jimma, south western Ethiopia. East African journal of public health. 2009;6(3).

61. Mengistu M, Shimelis T, Torben W, Terefe A, Kassa T, Hailu A. Human intestinal schistosomiasis in communities living near three rivers of Jimma town, south Western Ethiopia. Ethiopian journal of health sciences. 2011;21(2):111–8.

62. Mekonnen Z, Meka S, Zeynudin A, Suleman S. Schistosoma mansoni infection and undernutrition among school age children in Fincha’a sugar estate, rural part of West Ethiopia. BMC research notes. 2014;7(1):763.

63. Dufera M, Petros B, Erko B, Berhe N, Gundersen SG. Schistosoma mansoni infection in finchaa sugar estate: public health problem assessment based on clinical records and parasitological surveys, western Ethiopia. Science, Technology and Arts Research Journal. 2014;3(2):155–61.

64. Degarege A, Erko B. Prevalence of intestinal parasitic infections among children under five years of age with emphasis on Schistosoma mansoni in Wonji Shoa Sugar Estate, Ethiopia. PloS one. 2014;9(10):e109793.

65. Beyene G, Tasew H. Prevalence of intestinal parasite, Shigella and Salmonella species among diarrheal children in Jimma health center, Jimma southwest Ethiopia: a cross sectional study. Annals of clinical microbiology and antimicrobials. 2014;13(1):10.

66. Tulu B, Taye S, Amsalu E. Prevalence and its associated risk factors of intestinal parasitic infections among Yadot primary school children of South Eastern Ethiopia: a cross-sectional study. BMC research notes. 2014;7(1):848.

67. Hailu T, Yimer M. Prevalence of Schistosoma mansoni and geo-helminthic infections among patients examined at Workemeda Health Center, Northwest Ethiopia. Journal of Parasitology and Vector Biology. 2014;6(5):75–9.

68. Yimer M, Abera B, Mulu W. Scientia Research Library ISSN 2348-0416. Journal of Applied Science And Research. 2014;2(2):43–53.

69. Kure A, Mekonnen Z, Dana D, Bajiro M, Ayana M, Vercruysse J, et al. Comparison of individual and pooled stool samples for the assessment of intensity of Schistosoma mansoni and soil-transmitted helminth infections using the Kato-Katz technique. Parasites & vectors. 2015;8(1):489.

70. Jejaw A, Zemene E, Alemu Y, Mengistie Z. High prevalence of Schistosoma mansoni and other intestinal parasites among elementary school children in Southwest Ethiopia: a cross-sectional study. BMC public health. 2015;15(1):600.

71. Bajiro M, Dana D, Ayana M, Emana D, Mekonnen Z, Zawdie B, et al. Prevalence of Schistosoma mansoni infection and the therapeutic efficacy of praziquantel among school children in Manna District, Jimma Zone, southwest Ethiopia. Parasites & vectors. 2016;9(1):560.

72. Mekonnen Z, Suleman S, Biruksew A, Tefera T, Chelkeba L. Intestinal polyparasitism with special emphasis to soil-transmitted helminths among residents around Gilgel Gibe Dam, Southwest Ethiopia: a community based survey. BMC public health. 2016;16(1):1185.

73. Begna T, Solomon T, Yohannes Zenebe EA. Intestinal parasitic infections and nutritional status among primary school children in Delo-mena district, South Eastern Ethiopia. Iranian journal of parasitology. 2016;11(4):549.

74. Bajiro M, Dana D, Levecke B. Prevalence and intensity of Schistosoma mansoni infections among schoolchildren attending primary schools in an urban setting in Southwest, Ethiopia. BMC research notes. 2017;10(1):677.

75. Bajiro M GS, Hamba N, Alemu Y. Prevalence, Intensity of Infection and Associated Risk Factors for Schistosomamansoniand Soil Transmitted Helminthes among Two Primary School Children at nearby Rivers in Jimma Town, South West Ethiopia. Ann ClinPathol 2018;6(4):1144.

76. Teklemariam D, Legesse M, Degarege A, Liang S, Erko B. Schistosoma mansoni and other intestinal parasitic infections in schoolchildren and vervet monkeys in Lake Ziway area, Ethiopia. BMC research notes. 2018;11(1):146.

77. Mohammed J, Weldegebreal F, Teklemariam Z, Mitiku H. Clinico-epidemiology, malacology and community awareness of Schistosoma mansoni in Haradenaba and Dertoramis kebeles in Bedeno district, eastern Ethiopia. SAGE Open Medicine. 2018;6:2050312118786748.

78. Kebede T, Negash Y, Erko B. Schistosoma mansoni infection in human and nonhuman primates in selected areas of Oromia Regional State, Ethiopia. Journal of vector borne diseases. 2018;55(2):116.

79. Asfaw ST, Giotom L. Malnutrition and enteric parasitoses among under-five children in Aynalem Village, Tigray. Ethiopian Journal of Health Development. 2000;14(1):67–75.

80. Assefa A, Dejenie T, Tomass Z. Infection prevalence of Schistosoma mansoni and associated risk factors among schoolchildren in suburbs of Mekelle city, Tigray, Northern Ethiopia. Momona Ethiopian Journal of Science. 2013;5(1):174–88.

81. Mahmud MA, Spigt M, Mulugeta Bezabih A, Lopez Pavon I, Dinant G-J, Blanco Velasco R. Risk factors for intestinal parasitosis, anaemia, and malnutrition among school children in Ethiopia. Pathogens and global health. 2013;107(2):58–65.

82. Desta H, Bugssa G, Demtsu B. The Current Status of Schistosoma mansoni Infection among School Children around Hizaty Wedicheber Microdam in Merebmieti, Ethiopia. Journal of Bacteriology & Parasitology. 2014;5(5):1.

83. Abebe N, Erko B, Medhin G, Berhe N. Clinico-epidemiological study of Schistosomia mansoni in Waja-Timuga, District of Alamata, northern Ethiopia. Parasites & vectors. 2014;7(1):158.

84. Gebreegziabiher D, Desta K, Howe R, Abebe M. Helminth infection increases the probability of indeterminate quantiferon gold in tube results in pregnant women. BioMed research international. 2014;2014.

85. Alemu M, Kinfe B, Tadesse D, Mulu W, Hailu T, Yizengaw E. Intestinal parasitosis and anaemia among patients in a health center, North Ethiopia. BMC research notes. 2017;10(1):632.

86. Teshale T, Belay S, Tadesse D, Awala A, Teklay G. Prevalence of intestinal helminths and associated factors among school children of Medebay Zana wereda; North Western Tigray, Ethiopia 2017. BMC research notes. 2018;11(1):444.

87. Gebreyohanns A, Legese MH, Wolde M, Leta G, Tasew G. Prevalence of intestinal parasites versus knowledge, attitude and practices (KAPs) with special emphasis to Schistosoma mansoni among individuals who have river water contact in Addiremets town, Western Tigray, Ethiopia. PloS one. 2018;13(9):e0204259.

88. Nyantekyi LA, Legesse M, Belay M, Tadesse K, Manaye K, Macias C, et al. Intestinal parasitic infections among under-five children and maternal awareness about the infections in Shesha Kekele, Wondo Genet, Southern Ethiopia. Ethiopian Journal of Health Development. 2010;24(3).

89. Terefe A, Shimelis T, Mengistu M, Hailu A, Erko B. Schistosomia mansoni and soil-transmitted helminthiasis in Bushulo village, southern Ethiopia. Ethiopian Journal of Health Development. 2011;25(1):46–50.

90. Erko B, Degarege A, Tadesse K, Mathiwos A, Legesse M. Efficacy and side effects of praziquantel in the treatment of Schistosomia mansoni in schoolchildren in Shesha Kekele Elementary School, Wondo Genet, Southern Ethiopia. Asian Pacific journal of tropical biomedicine. 2012;2(3):235.

91. Wegayehu T, Tsalla T, Seifu B, Teklu T. Prevalence of intestinal parasitic infections among highland and lowland dwellers in Gamo area, South Ethiopia. BMC public health. 2013;13(1):151.

92. Mulu A, Legesse M, Erko B, Belyhun Y, Nugussie D, Shimelis T, et al. Epidemiological and clinical correlates of malaria-helminth co-infections in Southern Ethiopia. Malaria journal. 2013;12(1):227.

93. Degarege A, Animut A, Medhin G, Legesse M, Erko B. The association between multiple intestinal helminth infections and blood group, anaemia and nutritional status in human populations from Dore Bafeno, southern Ethiopia. Journal of helminthology. 2014;88(2):152–9.

94. Alemayehu B, Tomass Z. Schistosoma mansoni infection prevalence and associated risk factors among schoolchildren in Demba Girara, Damot Woide District of Wolaita Zone, Southern Ethiopia. Asian Pacific journal of tropical medicine. 2015;8(6):457–63.

95. Alemayehu B, Tomass Z, Wadilo F, Leja D, Liang S, Erko B. Epidemiology of intestinal helminthiasis among school children with emphasis on Schistosoma mansoni infection in Wolaita zone, Southern Ethiopia. BMC public health. 2017;17(1):587.

96. Tadege B, Shimelis T. Infections with Schistosoma mansoni and geohelminths among school children dwelling along the shore of the Lake Hawassa, southern Ethiopia. PloS one. 2017;12(7):e0181547.

97. Grimes JE, Tadesse G, Gardiner IA, Yard E, Wuletaw Y, Templeton MR, et al. Sanitation, hookworm, anemia, stunting, and wasting in primary school children in southern Ethiopia: Baseline results from a study in 30 schools. PLoS neglected tropical diseases. 2017;11(10):e0005948.

98. Tadesse G. The prevalence of intestinal helminthic infections and associated risk factors among school children in Babile town, eastern Ethiopia. Ethiopian Journal of Health Development. 2005;19(2):140–7.

99. Teklemariam Z, Abate D, Mitiku H, Dessie Y. Prevalence of intestinal parasitic infection among HIV positive persons who are naive and on antiretroviral treatment in Hiwot Fana Specialized University Hospital, Eastern Ethiopia. ISRN AIDS. 2013;2013.

100. Negussu N, Wali M, Ejigu M, Debebe F, Aden S, Abdi R, et al. Prevalence and distribution of schistosomiasis in Afder and Gode zone of Somali region, Ethiopia. Journal of global infectious diseases. 2013;5(4):149.

101. Schistosomiasis WECotCo, Organization WH. Prevention and control of schistosomiasis and soil-transmitted helminthiasis: report of a WHO expert committee: WHO; 2002.

102. Casavechia MTG, de Melo GdAN, Fernandes ACBDS, De Castro KR, Pedroso RB, Santos TDS, et al. Systematic review and meta-analysis on Schistosoma mansoni infection prevalence, and associated risk factors in Brazil. Parasitology. 2018:1–15.

103. Asundi A, Beliavsky A, Jian LX, Akaberi A, Schwarzer G, Bisoffi Z, et al. Prevalence of Strongyloides and Schistosomiasis among migrants: A systematic review and meta-analysis. International Journal of Infectious Diseases. 2018;73:228.

104. WHO. World health Organization (2010) Weekly epidemiological record. 2010;82:157–64.

105. Mas-Coma S, Valero MA, Bargues MD. Climate change effects on trematodiases, with emphasis on zoonotic fascioliasis and schistosomiasis. Veterinary parasitology. 2009;163(4):264–80.

106. Gali B, Nggada H, Eni E. Schistosomiasis of the appendix in Maiduguri. Tropical doctor. 2006;36(3):162–3.

107. Madinga J, Linsuke S, Mpabanzi L, Meurs L, Kanobana K, Speybroeck N, et al. Schistosomiasis in the Democratic Republic of Congo: a literature review. Parasites & vectors. 2015;8(1):601.

108. Moges, F., et al. Intestinal parasite infections in association with cutaneous fungal infection and nutritional status among schoolchildren in Tseda, northwest Ethiopia. Ethiopian Journal Health and Biomed Science, 2010, 3.1: 35–43

109. Alemu, Abebe, et al. Soil transmitted helminths and Schistosoma mansoni infections among school children in Zarima town, northwest Ethiopia. BMC infectious diseases, 2011, 11.1: 189.

110. WHO. Expert committe on the control of Schistosomiasis; World Health Organization. Prevention and control of schistosomiasis and soil-transmitted helminthiasis: report of a WHO expert committee. Who, 2002.

111. Abdulkadir, A., et al. Prevalence of urinary schistosomiasis in Nigeria, 1994-2015: Systematic review and meta-analysis. African Journal of Urology, 2017, 23.4.

